# Disentangling the spinal mechanisms of illusory heat and burning sensations in the Thermal Grill Illusion

**DOI:** 10.1101/2023.08.24.554485

**Authors:** Alexandra G. Mitchell, Jesper Fischer Ehmsen, Daniel Elmstrøm Christensen, Anna Villaume Stuckert, Patrick Haggard, Francesca Fardo

## Abstract

The Thermal Grill Illusion (TGI), a phenomenon in which the juxtaposition of innocuous warm and cold temperatures on the skin elicits a burning sensation, offers a unique perspective to how pain occurs in response to harmless stimuli. We investigated the role of the spinal cord in the generation of the TGI across two experiments (total n = 80). We applied heat and cold stimuli to dermatomes, areas of skin innervated by a single spinal nerve, that mapped onto adjacent or nonadjacent spinal segments. Enhanced warm and burning ratings during the TGI were observed when cold and warm stimuli were confined within the same dermatome. Further, we found the spatial organisation of warm and cold stimuli within and across dermatomes affected TGI perception. Perceived warmth and burning intensity increased when the cold stimulus projected to the segment more caudal to the warm stimulus, whilst perceived cold during the TGI decreased, compared to the opposite spatial arrangement. This suggests the perception of TGI is enhanced when cold afferents are projected to spinal segments positioned caudally in relation to those receiving warm afferents. Our results indicate distinct interaction of sensory pathways based on the segmental arrangement of afferent fibres and are consistent with current interpretations of the spread and integration of thermosensory information along the spinal cord.

## Introduction

The thermal grill illusion (TGI) is the sensation of burning heat or pain when harmless cold and warm temperatures are simultaneously applied to the skin [5,6]. As the cold and warm temperatures are innocuous and therefore insufficient to activate peripheral nociceptors, the generation of illusory heat is thus attributed to central nervous system mechanisms [5,8,9,11].

The TGI is often described as encompassing two distinct perceptual components - an illusion of heat and an illusion of pain [7,8]. Historically, the thermosensory and painful components of the TGI were explained by distinct spinal and supraspinal mechanisms, respectively [5]. The illusory pain component has been ascribed to a disinhibition mechanism at the level of the thalamus, primarily based on the observations of unremitting pain following thalamic lesions [4,5]. Some more recent human studies on TGI have posited that the illusory pain component of the TGI depends uniquely on supraspinal mechanisms, based on the observed modulation of the illusion in accordance with a spatiotopic rather than somatotopic representation of the body [17] and that the experience of TGI is not modulated by tactile gating - a spinally mediated process involving inhibition of nociceptive activity by concurrent somatosensory activity [10].

Counter to this perspective, other research has found the TGI varies depending on whether cold and warm afferents mapped either onto adjacent or non-adjacent spinal segments [9]. This suggests that the spinal cord is an initial site of thermosensory integration underlying the TGI. Further support for spinal mechanisms comes from research demonstrating that noxious heat and the TGI were comparably reduced by conditioned pain modulation in humans, a mechanism that is mediated by descending modulatory systems that originate in the brain but acts on the spinal cord [11]. These findings collectively refute the notion of a purely supraspinal hypothesis of TGI and underscore the significance of spinal mechanisms in the manifestation of both illusory heat and pain.

In this paper our objective was to directly investigate the hypothesis that thermosensory and burning components of the TGI are mediated by spinal mechanisms in humans. Cold and warm stimuli were presented at a fixed distance on the skin but depending on their orientation on the arm, they elicited differing neural activity in the spinal cord (Figure 1 A and B). Our assumption was that cold and warm-related neural activity would show stronger integration, and thus stronger TGI effects, when the stimuli mapped to the same or adjacent spinal segments, than when they mapped onto segments that were anatomically further apart. Additionally, we investigated spatial order effects associated with the integration of cold and warm sensory information at the dermatome (skin) and segmental (spine) levels (Figure 1B). If the experience of the TGI is influenced by the relative location of warm and cold afferents in different spinal segments, this finding would provide compelling, additional evidence that the spine serves as an initial site for the cold-warm integration that produces perceptions of illusory heat and pain.

**Figure 1:**
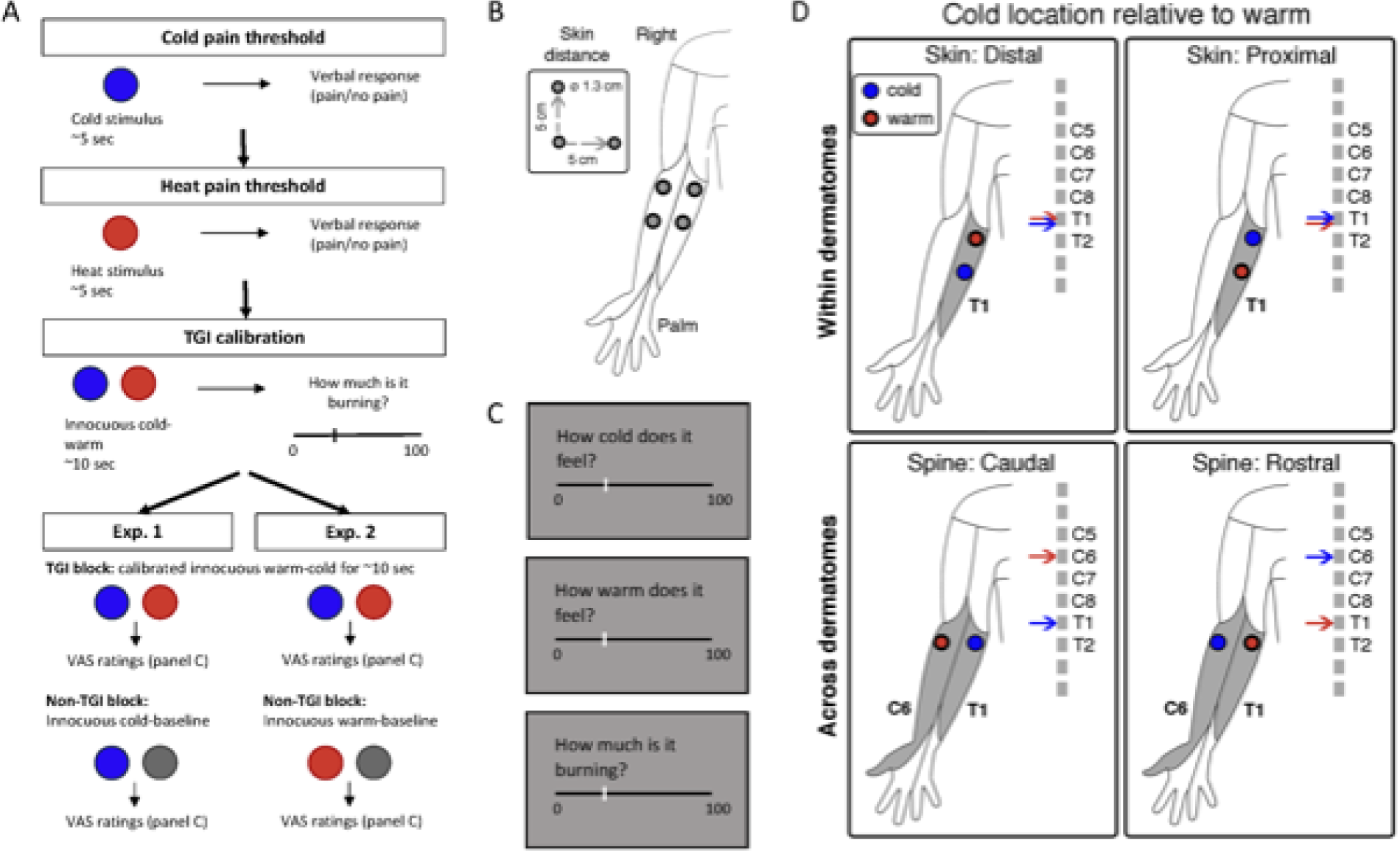
(A) An example sequence of an experiment. The order of pain threshold blocks 1 and 2 and experimental sessions 1 and 2 were counterbalanced evenly across participants. (B) Placement and distance of thermodes on the inner forearm. (C) The three Visual Analogue Scales (cold, warm, burning) participants used to report sensation coming from the reference probe, presented in a randomised order for every trial. (D) The corresponding spinal mapping of the probe placement for all four conditions. Within dermatomes, the relative location of the cold thermode was proximal or distal. Across dermatomes, the relative location of the corresponding spinal segments was rostral or caudal. Within dermatome conditions also included warm and cold thermodes in C6 (not depicted here), as well as T1. The size of the stimulus site in panels C and D probe is not to scale.

## Methods

### Participants

The study entailed two separate experiments, collectively involving 80 healthy volunteers. Forty participants took part in experiment 1 (27 females and 13 males, mean age = 25.38 years old, SD = 4.67, range = 18 - 36) and another 40 participants in experiment 2 (25 females and 14 males and 1 non-binary, mean age = 25.73 years old, SD = 4.12, range = 21 - 39). Data collection for experiments 1 and 2 took place between August - December 2022 and January - May 2023 respectively and there were no participants that took part in both experiments. The research methodology complied with the principles set forth in the Declaration of Helsinki and received ethical approval from the Institutional Review Board (IRB) at the Danish Neuroscience Center, Aarhus University, Denmark. Prior to commencing the study, all participants were fully informed about the procedures and provided their voluntary consent.

### Stimuli

All thermal stimuli were delivered using two NTE-3 Thermal Sensitivity Testers (PhysiTemp Instruments LLC, 10mm in diameter) controlled by PhysiTemp NTE-3 software (version 5.4b). The procedure involved measurements of heat and cold pain thresholds, calibration of cold-warm temperature pairs eliciting TGI, and an experimental task where TGI and non-TGI stimuli were applied on dermatomes that mapped onto adjacent or nonadjacent spinal segments. The thermodes were heated or cooled to the required stimulus temperature prior to being placed on the participant’s skin. If the skin temperature affected the surface temperature of the probe, the experimenter waited to start the trial until the desired temperature was reached. The maximum rate of temperature change of the probes was 1°C/s.

The Thermal Grill Illusion (TGI) is characterised by two key phenomena: thermosensory enhancement and illusory pain. Thermosensory enhancement refers to an amplified perception of heat or cold when cold and warm stimuli are simultaneously applied, as opposed to when each stimulus is presented individually or paired with a neutral temperature. Notably, the majority of individuals experience an intensification of heat rather than cold [8]. Illusory pain, on the other hand, denotes the perception of a burning sensation elicited by the pairing of warm and cold stimuli, an experience that is largely absent or significantly diminished when each stimulus is presented alone or combined with a neutral temperature. Thus, indicators of a stronger TGI are reduced cold ratings, coupled with heightened warm and burning ratings. The last of these is generally considered to be the most salient feature of the TGI, and has received the most attention within the TGI literature.

While most previous studies have focused on the TGI as a single experience, explicitly understood as an experience of or like pain, our approach differed. We instructed participants to report the sensation from just one of the thermal components, corresponding to the cold thermode location (Exp. 1) or the warm thermode location (Exp. 2). Importantly, the aim of this method was not to isolate any possible independent effect of warm and cold stimulation during TGI, but rather to ensure consistency in participants’ ratings across trials and different stimulus conditions.To investigate both the thermosensory and burning components in each experiment, we asked participants to provide a subjective evaluation of multiple sensory qualities - cold, warmth and burning (pain). Stimuli consisted of either cold-warm pairs (TGI stimuli), which potentially evoked an illusion of heat and pain, or non-TGI control stimuli. The non-TGI control stimuli were constructed by pairing the cold (Exp. 1) or the warm (Exp. 2) component of the TGI stimuli, with a baseline temperature of 30°C. All stimulation pairs were presented at a fixed distance on the skin, either within the same dermatome or across dermatomes that mapped onto non-adjacent spinal segments (Fig. 1). This arrangement of cold and warm probes on the skin was specifically chosen to be able to explore the spinal mechanisms of the TGI. Similar TGI stimulus designs based on just two thermal probes, have been used previously [7,9].

### Procedure

An outline of the procedure for each experiment is found in Figure 1A. We measured cold and heat pain thresholds in a stepwise manner, the order of which was counterbalanced across participants. For cold pain, a single thermode at a starting temperature of 25°C was held on the participant’s dorsal forearm for five seconds. After which, the participant verbally reported (yes/no) any experience of pain. If the participant reported no pain, the temperature of the thermode was lowered by 5°C, and placed back on the skin for another five seconds after which the participant reported whether they experienced pain. This step was repeated either until th participant responded ‘yes’ or the thermode reached the set minimum temperature of 5°C. If the participant responded ‘yes’ before the minimum temperature, the temperature of the thermode was increased by 1°C until the participant no longer experienced pain. The cold pain threshold was identified as the highest temperature at which the participant reported a painful experience. The same steps were repeated for heat pain, but with increasing intervals of 5°C and with a starting temperature of 35°C and a maximum temperature of 45°C. The heat pain threshold was identified as the lowest temperature at which the participant reported a painful experience. Heat and cold pain thresholding procedures were completed once per participant.

We calibrated TGI stimuli by identifying a cold-warm temperature pair based on specific criteria: (1) consistently eliciting a burning sensation of at least 15 on a scale ranging from 0 to 100, (2) consistently avoiding a burning sensation (less than 15) when the cold-neutral (Exp. 1) or warm-neutral (Exp. 2) stimuli were presented, (3) both cold and warm temperatures falling within the innocuous range based on individual cold and heat pain thresholds. TGI temperature pairs, starting at 25°C and 35°C were presented on the participant’s forearm for 10 seconds. After 10 seconds, the participant had to rate the perceived burning (using a VAS scale from 0 - 100) coming from either the cold (Exp. 1) or warm Exp. 2) probe. If the participant did not rate their perceived burning as above 15 on the scale, the experimenter increased the temperature on the warm probe, and decreased the temperature on the cold probe systematically. They then placed the probes at a different location on the forearm and repeated the trial. If the participant rated above 15 on the VAS, the temperature combination was repeated another six times. To determine the TGI temperatures, the participant needed to rate the probe as above 15 on the burning VAS four times out of six. In situations where the participant did not consistently report burning for temperatures that were below their pain thresholds, the maximum and minimum possible temperatures were used (2°C below their thresholds). For pain threshold measurements and TGI calibration, we positioned the probes within a single dermatome.

To address our experimental questions, we presented the calibrated TGI stimuli, as well as cold-neutral (Exp. 1) or warm-neutral (Exp. 2) non-TGI stimuli using two thermodes. In non-TGI stimuli the cold or warm temperatures were set to match the temperature used for TGI stimulation, but paired with a neutral temperature set at 30°C.

The two thermodes were positioned on the internal surface of either forearm, with a constant spacing of 5 cm in each direction (Fig. 1B). In rare cases where the participant’s forearm was too narrow to position the probes at 5 cm apart across dermatomes, this distance was adjusted either 4.5 or 4 cm. The positioning of the thermodes was either within the same dermatome (C6 and T1) or across dermatomes mapped onto non-adjacent spinal segments (i.e. C6 - T1). Further, we manipulated the spatial arrangement of the temperature pairs, by systematically presenting an equal number of trials where the cold thermode was applied on a proximal or distal location within a dermatome, or was applied on a dermatome that mapped onto a rostral or caudal segment along the spinal cord. We based the demarcation of the dermatome boundaries on the American Spinal Injury Association (ASIA) map and positioned the thermodes in relation to standard anatomical landmarks. Proximo-distal coordinates referred to locations on the skin closer to the elbow or the wrist, whereas rostral-caudal coordinates referred to spinal segments closer to the head (C6) or the lower back (T1). Spatial arrangements of stimuli are depicted in Figure 1D. The order of the stimuli (TGI vs. non-TGI), the dermatome condition (within vs. across) and the relative placement of the colder temperature (proximal vs. distal or rostral vs. caudal) were pseudo-randomised and counterbalanced between participants.

During each trial, the experimenter positioned the two thermodes on the participant’s skin, and then waited for both thermodes to reach within .25°C of the desired temperatures for each stimulus. After ten seconds of stimulation, participants reported their ratings using three sequential VAS scales for each perceptual quality (cold, warm and burning). Scales ranged from 0, indicating the lack of the corresponding sensory quality, to 100, indicating an extreme sensation (Fig. 1C) and were presented in a random order. The order of the three VAS scales was randomised across trials and participants had a maximum of eight seconds to respond to each scale. For each scale, participants provided their responses using the arrow keys on a keyboard and rated the intensity of their sensations from a specific location (labeled ‘A’ or ‘B’), based on the experimenter’s instruction. Unbeknown to the participant, this location systematically corresponded to either the colder temperature (Exp. 1) or the warmer temperature (Exp. 2). An auditory cue (300Hz, 100ms) indicated when the participants completed all ratings, after which the experimenter removed the thermodes from their skin. We presented a 200ms fixation dot before beginning the next trial. Each thermode configuration was tested three consecutive times on each arm, on three different and non-overlapping skin locations. An auditory tone of 500Hz lasting 100ms was played to indicate to the experimenter that the trial was over, and that they should rearrange the thermode configuration. Two simultaneous tones of the same quality occurred every three trials, indicating that the participant should change arms to stimulate different dermatomes depending on a pseudo-randomisation order. Tones were used because the experimenter could not see the screen, therefore it was the most effective way to inform the experimenter when each trial was completed. Each of the four experimental conditions was repeated 12 times, with both the right and left forearms stimulated, and a minimum of five trials between the re-stimulation of the same skin location. This method ensured that the same skin locations were not stimulated consecutively to minimise the potential of carry-over effects.

Experiments 1 and 2 were conducted in two independent groups of participants and followed exactly the same procedure except for two elements. In Experiment 1, participants rated the sensations localised underneath the colder thermode, and the non-TGI stimuli corresponded to cold-neutral pairs, where the temperature of the cold thermode in both conditions was the same. In Experiment 2, participants rated the sensations localised underneath the warmer thermode, and the non-TGI stimuli corresponded to warm-neutral pairs, where the temperature of the warm thermode in both conditions was the same.

### Sample size

An initial pilot study informed the pre-registered (https://osf.io/4xcn5/) calculation of the sample size. To test the directional TGI hypothesis with 95% power and detect an effect size for the coefficient of .12 or greater, we determined that we needed a minimum number of 32 TGI-responsive participants. We defined TGI-responders as those individuals for whom the median burning ratings for TGI stimuli significantly exceeded 0. Non-responders were individuals that did not meet this criterion when tested with the max cold-warm temperatures allowed in the experiment. The predefined cut-off for TGI stimulation was 10°C and 44°C, due to both limitations of the thermode and to reduce likelihood of sensitization to heat stimuli. In Experiment 1, recruitment continued until we achieved the target of 32 TGI-responsive participants. We verified this criterion every 10 participants, resulting in a total sample size of 40 participants with 32 TGI responders. In Experiment 2, we stopped recruitment once we collected data from 40 participants, which resulted in a total of 37 TGI responders. This decision was based on meeting both required criteria: (1) matching the sample size of Exp. 1 for consistency, and (2) achieving the minimum requirement of 32 TGI-responsive participants as determined by the power analysis.

### Data analyses

We re-scaled data from cold, warm and burning VAS ratings from their original values to a range of 0 to 1. Following re-scaling, we applied zero-inflated mixed-effects beta regression models separately for each set of VAS ratings. In these models, we incorporated three fixed effects; the type of stimulation (non-TGI vs. TGI), the dermatome condition (within the same dermatome vs. across different dermatomes) and the spatial positioning of the cold or neutral thermode (proximal vs. distal within dermatomes; rostral vs. caudal across dermatomes). This allowed us to assess the individual and interactive effects of these three factors on VAS ratings. Further, we added random intercepts of subject and trial order to our models to account for between-subject variability and the effects of repeated measures.

The choices of the zero-inflated approach and the use of beta regressions were necessitated by the specific distribution of VAS ratings. The beta distribution is suitable for modelling VAS rating data, as they are proportional in nature. Additionally, the zero-inflation was needed due to the presence of an excess number of zero values in specific ratings and conditions. Specifically, we anticipated an over-representation of zero values for thermosensory ratings that were counterfactual to the objective stimulation quality (i.e., cold ratings of warm stimuli and warm ratings of cold stimuli) and burning ratings of non-TGI stimuli. The latter stimuli were designed to not elicit an illusion or trigger a weaker illusion as compared to the TGI stimuli. We carried out the statistical analyses using the ‘glmmTMB’ package in R (version 1.1.8), and statistical significance was set at p < .05.

For data presentation purposes, the median VAS ratings for each sensory quality (cold, warm, burning) were calculated per simulation type (TGI, non-TGI), dermatome (within, across) and spatial location (proximo-distal, rostro-caudal) for each participant. For hypothesis one, these values were further averaged so there were only four values per participant, for each sensory quality (within and across dermatome ratings for TGI and non-TGI stimuli). For analysis of burning VAS ratings, only those participants who were deemed as TGI responders (n = 32 for Exp. 1 and n = 37 for Exp. 2) were included.

### Data availability

The experimental procedure, power analyses to determine sample size and statistical approach were preregistered for both experiment 1 (https://osf.io/4xcn5/) and experiment 2 (https://osf.io/dhg8u/).

All raw data and code for the analysis are available in the GitHub repository (https://github.com/Body-Pain-Perception-Lab/tgi-spinal). This, and a wiki guide to analysing the data, can be accessed through the project’s OSF page (https://osf.io/uyrtq/).

## Results

The full results for both experiments 1 and 2 are summarised in Figure 2.

**Figure 2:**
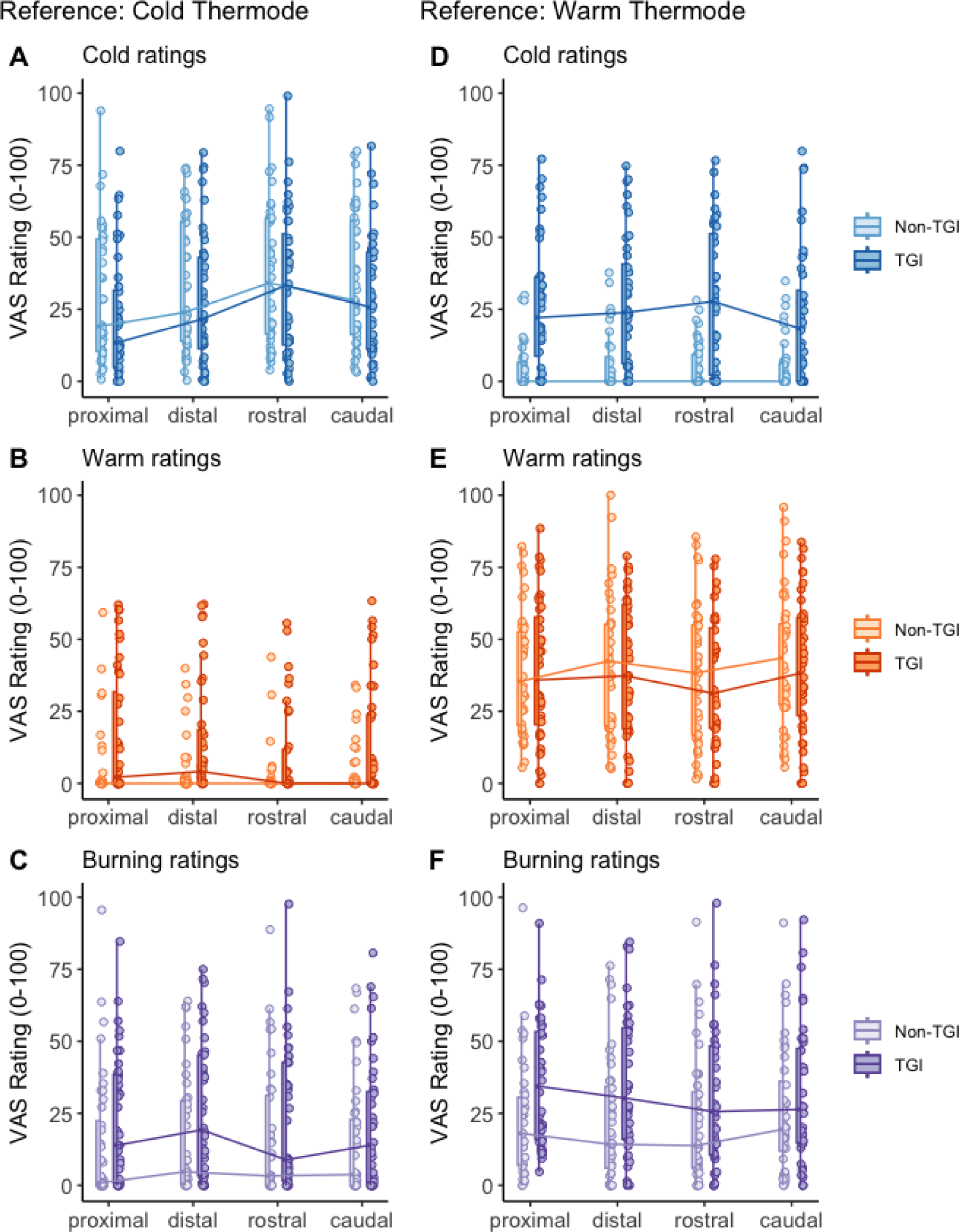
Individual median VAS ratings for Experiment 1 (A-C) and Experiment 2 (D-F) across stimulus manipulation, and spatial location. Proximal and distal locations are within dermatome, rostral and caudal locations are across dermatomes. All spatial locations refer to the location of the cold thermode, compared to the warm. Box plots show median and interquartile range. For completion, the data presented here include trials where VAS ratings equal 0, which are modelled seperately in the main analyses.

### Thermosensory and burning components of TGI perception are spinally mediated

The typical heat and burning perception associated with TGI was more robust when stimuli were confined within dermatomes compared to when applied across dermatomes, corresponding to non-adjacent spinal segments. When rating the cold thermode (Exp. 1, Fig. 3A), participants reported a stronger reduction in the subjective experience of cold specifically for TGI stimuli applied within a dermatome compared to across dermatomes. The results of the zero-inflated beta regression show a stimulation by dermatome interaction (cold ratings: *β* = -.15, p < .01), alongside an increased subjective experience of warmth for both TGI and non-TGI stimuli (dermatome main effect: *β* = .26, p < .001; stimulation by dermatome interaction: *β* = -.03, p = .77). Further, participants reported no significant modulation of burning ratings depending on the dermatome condition (stimulation by dermatome interaction: *β* = .10, p = .15; dermatome main effect: *β* = -.04, p = .39). These results indicated that when participants judged the cold thermode, the greatest modulation in TGI perception was related to cold perception, with within-dermatome TGI stimuli perceived as the least cold. While the modulation of cold perception was specific for TGI, increased warmth was reported irrespective of whether the cold thermode was paired with a warm (TGI stimuli) or neutral temperature (non-TGI stimuli) within a dermatome. Overall, these findings are in line with the notion that TGI can be considered a misperception of cold [8,9].

**Figure 3:**
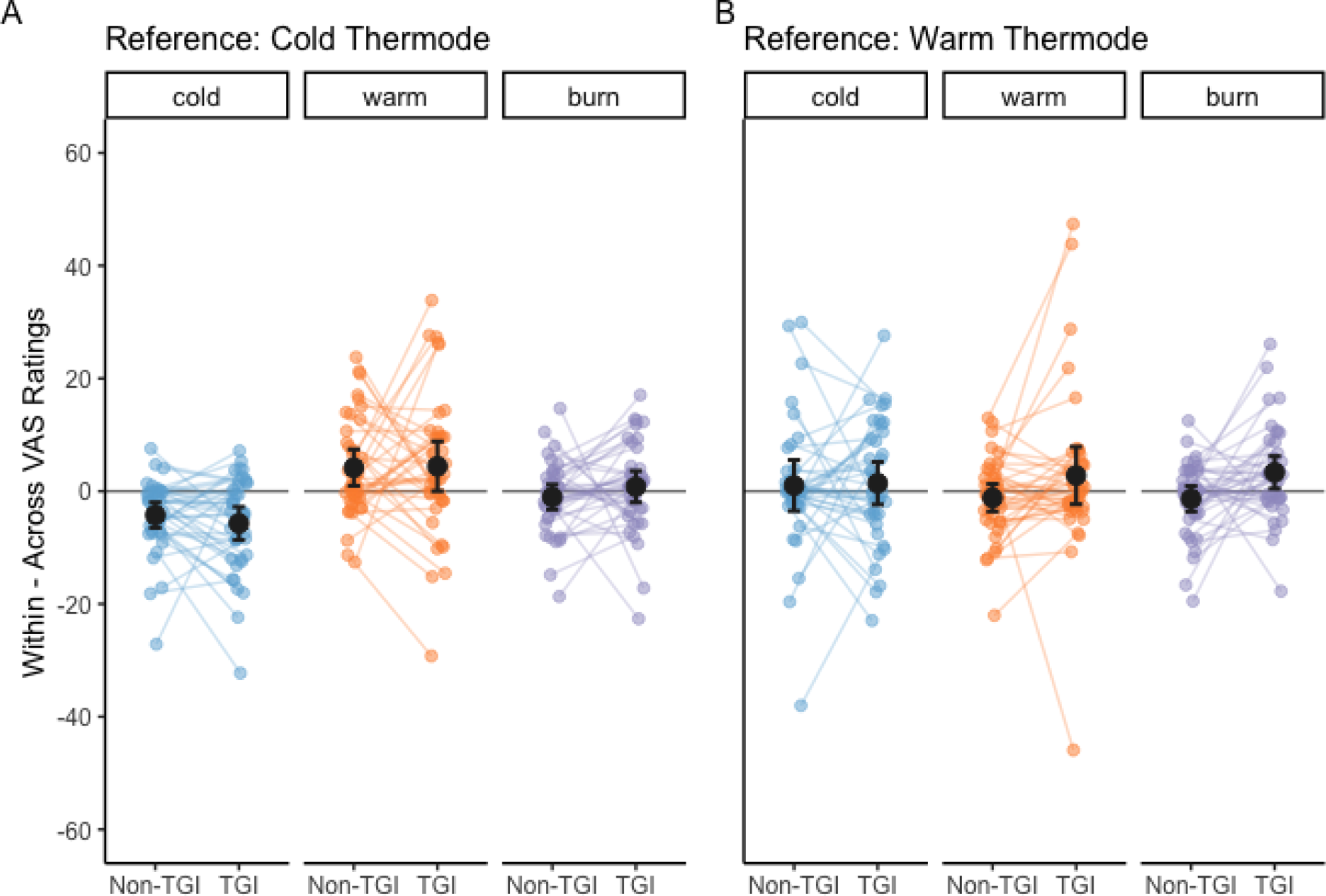
Thermosensory and burning components of TGI are spinally mediated: The placement of warm and cold stimuli in different dermatomes reduces the experience of the TGI. The difference between within and across dermatome conditions are displayed for each type of stimulation (Non-TGI and TGI) for each VAS rating quality (cold, warm and burning). Positive values represent higher ratings within dermatomes, negative values represent higher ratings across dermatomes. In experiment one (A), where participants judged sensations at the location of the cold thermode, cold ratings were significantly reduced during TGI when thermodes were placed across dermatomes. In experiment two (B), where participants judged sensations at the location of the warm thermode, warm and burning ratings were significantly reduced during TGI. Small dots are individual subject means, large dots are population means for each condition and error bars are 95% confidence intervals. Note that to best reflect the outcome of our zero-inflated statistical model this figure only includes trial data where VAS ratings were greater than 0.

When rating the warm thermode (Exp. 2, Fig. 3B), participants reported markedly enhanced burning sensations for TGI, but not non-TGI, stimuli applied within a dermatome compared to across dermatomes (dermatome main effect: *β* = -.07, p = .11; stimulation by dermatome interaction: *β* = .21, p < .001). However, we did not observe modulation of cold (dermatome main effect: *β* = .01, p = .83; stimulation by dermatome interaction: *β* = .08, p = .28) or warm ratings by dermatome (dermatome main effect: *β* = -.04, p = .28; stimulation by dermatome interaction: *β* = .07, p = .16). Overall, these results indicated that when participants judged the warm thermode, the greatest modulation in TGI perception was related to burning sensations, with within-dermatome TGI stimuli perceived as the most burning.

### Proximodistal bias in cold perception

Previous research has demonstrated a phenomenon known as distal inhibition which occurs when two heat stimuli are presented simultaneously. Typically, heat pain ratings at the proximal location are lower, and therefore inhibited, when the temperature at the distal location also produces heat pain, compared to when it is neutral [19,20]. When applied to our study, distal inhibition would result in higher ratings associated with both burning and the congruent sensation when the reference thermode (cold for Exp. 1, warm for Exp. 2) is in a more distal location compared to the other thermode. We tested the occurrence of this distal inhibition effect to the perception of mild temperatures in TGI and non-TGI stimuli within single dermatomes (Fig. 4).

**Figure 4:**
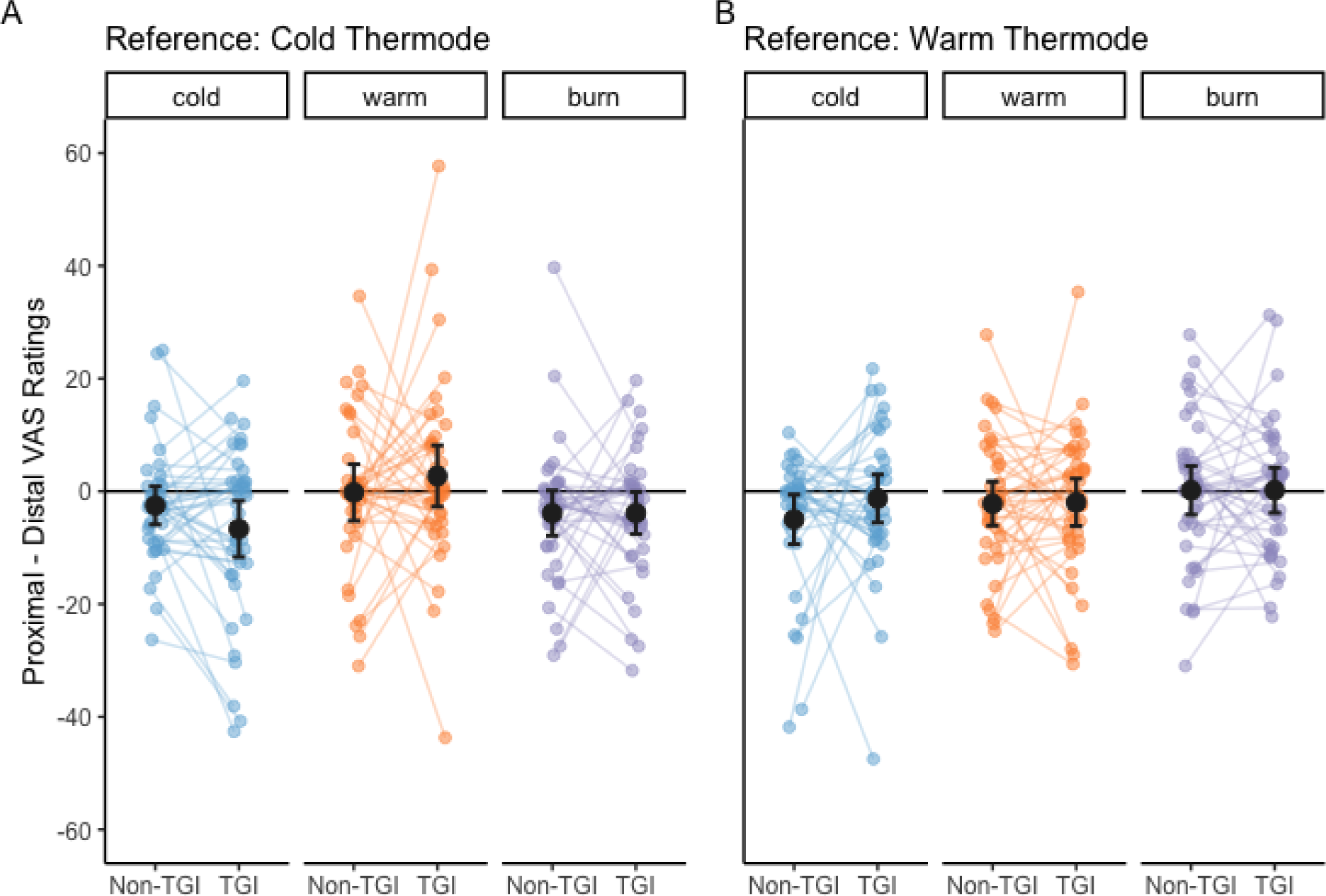
Proximodistal bias in cold perception: The spatial location of the warm and cold thermodes within dermatomes affects the perception of cold. The difference between the proximal and distal location of the reference thermode in the within dermatome are displayed by condition and by stimulation type (Non-TGI, TGI) for all VAS rating types (cold, warm, burn). Positive values show higher ratings when the cold probe is more proximal than the warm probe. In experiment one (A), where participants judged sensations at the location of the cold thermode, cold and burning ratings were higher when the cold probe was more distal. In experiment two (B), where participants judged sensations at the location of the warm thermode, cold ratings were higher when the cold probe was more distal. Small points show data from each participant, large dots are means across trials for each condition, and error bars show 95% confidence intervals. Note that to best reflect the outcome of our zero-inflated statistical model this figure only includes trial data where VAS ratings were greater than 0.

We found that cold and burn perception were modulated by the proximodistal location of the cold thermode. In experiment 1, cold ratings (dermatome main effect: *β* = -.18, p < .001; stimulation by dermatome interaction: *β* = -.08, p = .30) and burn ratings (dermatome main effect: *β* = -.17, p < .05; stimulation by dermatome interaction: *β* = -.09, p = .37) were enhanced when the reference (cold) thermode was located more distally than the warm probe, irrespective of whether the stimulus was TGI or non-TGI (Fig. 4A). We found a similar finding for cold perception in experiment 2 (dermatome main effect: *β* = -.18, p < .05; stimulation by dermatome interaction: *β* = .10, p = .34), where cold ratings were higher when the reference (warm) thermode was more proximal than the cold thermode (Fig. 4B). These findings suggest that the notion of distal inhibition can be extended to innocuous cold perception and burning sensations that are not specific to TGI at objectively mild temperatures.

### Directional effects in inter-segmental sensory integration

A main objective of this study was assessing spatial order effects along the rostrocaudal axis at the spinal level. In the across dermatome condition, we delivered an equal number of trials in which the cold stimulus was applied on a dermatome that mapped more rostrally or caudally compared to the warm. We found thermosensory components of the TGI were enhanced when cold sensory afferents mapped on to more caudal spinal segments, compared to warm sensory afferents.

In experiment 1, the modulation of thermosensory ratings corresponded to significant rostrocaudal main effects for both cold ratings (*β* = -.15, p < .01) and warm ratings (*β* = .21, p < .05), but this effect was not specific for TGI stimuli (stimulation by rostrocaudal location interaction, cold ratings: *β* = -.10, p = .19; warm ratings: *β* = .06, p = .65). In experiment 2, the modulation of cold ratings was specific for TGI stimuli (stimulation by rostrocaudal location interaction: *β* = -.23, p < .05), while the modulation of warm ratings was significant irrespective of stimulation type (rostrocaudal main effect: *β* = .19, p < .001; stimulation by rostrocaudal interaction: *β* = .01, p = .94). The rostrocaudal mapping of cold-related activity also modulated burning ratings irrespective of stimulation type, in experiment 2 (rostrocaudal main effect: *β* = .17, p < .01; stimulation by rostrocaudal location: *β* = -.14, p = .10).

Therefore, when rating either the cold thermode (Exp. 1, Fig. 5A) or the warm thermode (Exp.2, Fig. 5B), participants reported a reduced subjective experience of cold, alongside an enhanced perception of warmth when the cold thermode was applied on a dermatome that mapped onto a more caudal segment.

**Figure 5:**
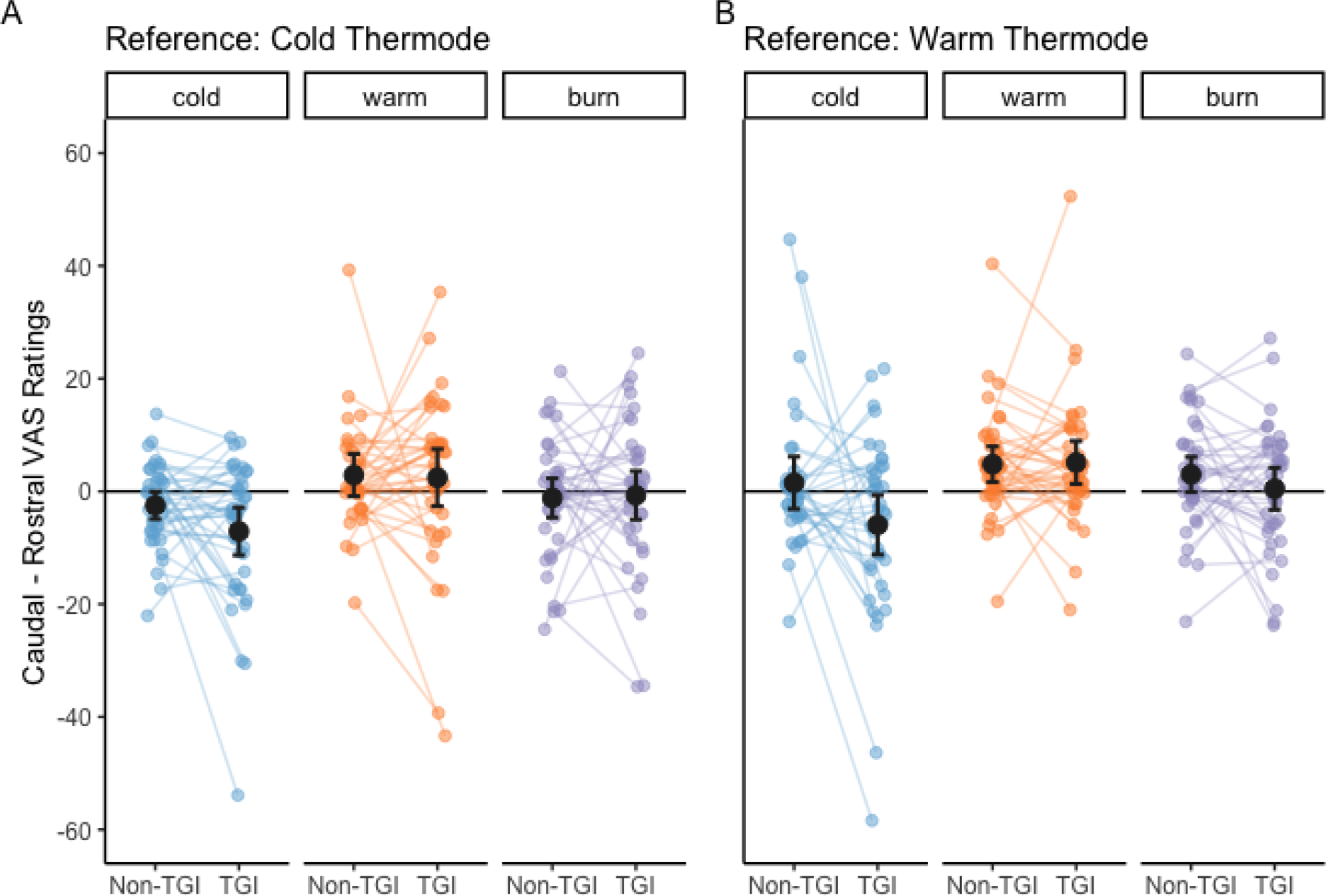
Directional effects of inter-segmental sensory integration: Spatial location of warm and cold thermodes across dermatomes affects the perception of TGI and non-TGI stimuli. The difference between the caudal and rostral dermatomes are displayed by conditions and stimulation type (Non-TGI, TGI) for all VAS rating types (cold, warm, burn). Positive values represent higher ratings when the cold probe is located in dermatomes that are related to more caudal spinal segments (T1) than the warm probe (C6). In experiment one (A), where participants judged sensations at the location of the cold thermode, perception of warm and cold, cold ratings were reduced and burning ratings increased when the cold probe was located in more caudal dermatomes. In experiment two (B), where participants judged sensations at the location of the warm thermode, the changes in warm and cold ratings were similar to experiment one but specific to TGI stimuli. Small points show data from each participant, large dots are means across trials for each condition, and error bars show 95% confidence intervals. Note that to best reflect the outcome of our zero-inflated statistical model this figure only includes trial data where VAS ratings were greater than 0.

## Discussion

In this study, we showed that organisation of cold and warm primary afferents both within and across segmental locations affects perception of the TGI. Our findings suggest that the thermosensory quality of the reference thermode (cold or warm) showed differential sensitivity to thermosensory and painful aspects of the TGI experience, and collectively suggested that both qualitative components of the illusion are modulated at the spinal cord level. This interpretation is consistent with a previous study using a similar dermatome manipulation [9], as well as another study showing modulation of heat and pain ratings of TGI stimuli by conditioned pain modulation [11]. In addition to this, we found a notably enhanced TGI effect when the cold stimulus induced more caudal activity within the spinal cord.

Our TGI stimuli did not use the traditional arrangement of alternating warm and cold bars in our study are not the conventional arrangement, however, research has shown that the TGI can be induced from a variety of alternating warm-cold patterns on the skin [15]. In addition, the stimuli used in our paradigm have been previously established to produce TGI [7,9]. Overall these results support the role of spinal processes in the generation of distinct perceptual aspects of the TGI and are in line with the interpretation that the thermosensory and burning components of the TGI can differ strongly according to the location of activated cold-warm primary sensory afferents in the spinal cord. They also highlight the importance of the veridical temperature of the stimulus being judged when assessing thermosensory and burning components of TGI perception.

The details of spinal neuroanatomy provide insightful perspectives concerning the two main findings of these experiments (Fig.6): (1) reduced cold, enhanced heat and burning sensations when thermosensory integration takes place more focally within a few spinal segments, and (2) clear effects when distinct cold and warm stimuli elicited a differential spatial pattern of neural activity along several spinal cord segments.

**Figure 6:**
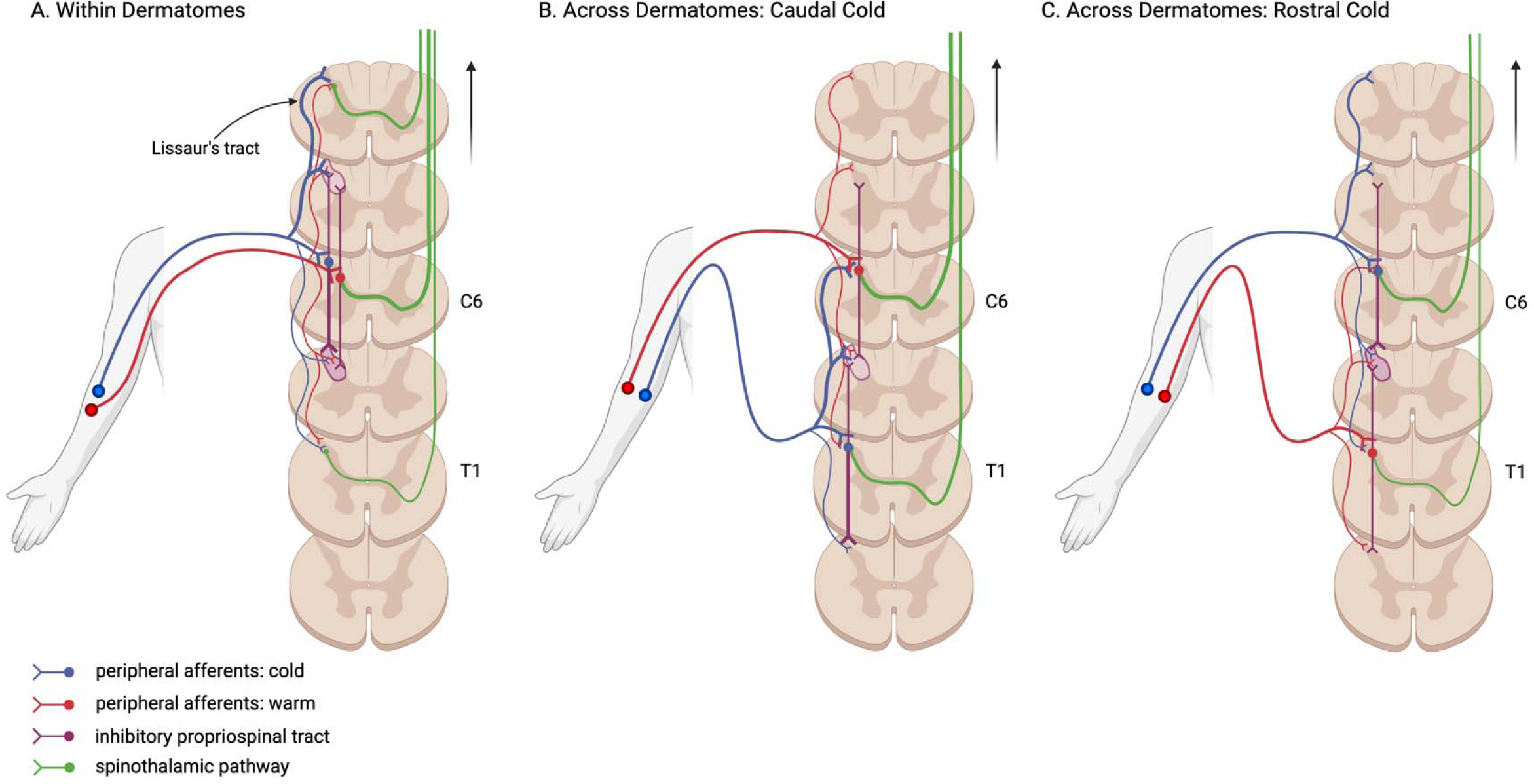
Schematic of the spinal neuroanatomy associated with warm and cold afferents (A) within the same spinal segment and across two separate non-adjacent spinal segments where (B) cold afferents project to more caudal segments and (C) cold afferents project to more rostral segments. After entering the spinal cord, peripheral warm and cold afferent fibres form collateral branches and synapses above and below the level of entry in neighbouring spinal segments. Propriospinal neurons possibly inhibit responses in neighbouring segments (i.e., inter-segmental inhibition), which means integration of warm and cold sensory signals is more likely within (A) than across (B and C) dermatomes. Our finding of stronger TGI when the cold afferents are in more caudal spinal segments (B) compared to rostral (C) could be due to an asymmetrical distribution of responding primary afferents along the Lissaur’s tract, or larger descending inhibitory propriospinal connections that are specific for cold. This predicted asymmetry is depicted here through larger/narrower line widths in both the Lissaur’s tract and propriospinal tract. Inhibition within neighbouring segments of the dorsal horn from propriospinal neurons is illustrated in purple, with segments where we expect a stronger inhibitory response represented by a darker shade, compared to where there is less inhibition. Projection neurons decussate and ascend contralaterally forming the spinothalamic pathway. They are shown here only in segments where cold and warm afferents can combine to produce a TGI sensation, and once again linewidth represents the predicted signal strength of combined warm and cold response. The black arrow indicates direction of supraspinal structures such as the thalamus.

Small primary afferents, responsible for mediating temperature and pain sensations, split into ascending and descending branches that cover one to two segments before they enter the dorsal horn [12,13]. This pattern forms the Lissauer’s tract, a structure hypothesised to regulate sensory transmission to the dorsal horn and influence spinal receptive field size [25]. Additionally, the endings of small primary afferents within the superficial laminae of the dorsal horn form synapses with both propriospinal neurons and projection neurons that target supraspinal structures known to significantly influence TGI perception [6,14,16]. Evidence from animal studies shows that propriospinal neurons, confined within the spinal cord, exhibit bidirectional collateral branches along the rostrocaudal plane [22,23]. These connections shape the network of interneurons that modulates sensory information delivered to the dorsal horn [18,24].

Our results indicate that enhanced TGI perception when cold-warm stimuli are applied within dermatomes (Fig. 6A), compared to across dermatomes (Fig. 6B) may be attributed to a confluence of interconnected mechanisms. First, the Lissauer’s tract, with its short rostrocaudal span of only one to two segments, aligns with the constraints of individual dermatome boundaries. This tract potentially facilitates the integration of warm and cold sensory information within the spinal cord, explaining reduced TGI percepts when cold-warm afferents span multiple spinal segments. Second, spinal circuits formed by propriospinal neurons may promote this sensory integration within a given spinal receptive field. They may do this by inhibiting activity in adjacent fields, a mechanism that corresponds to the principle of lateral inhibition which is present in both peripheral and central nervous systems and influences various sensory modalities, such as thermoception and nociception [1,2,21]. In the specific framework of the TGI, previous studies have suggested that TGI perception is related to the difference between cold and warm temperatures, with a greater difference leading to a higher likelihood of TGI perception [3]. If lateral inhibition is involved, enhanced TGI perception when cold-warm afferents are within dermatomes is expected, as the larger the difference, the greater the contrast and the greater the lateral inhibition. This understanding aligns with the potential role of lateral inhibition in accentuating the illusory sensations of heat and pain in the TGI by amplifying the differences between simultaneous cold and warm stimulation. Taken together, the spatial characteristics of the Lissauer’s tract, the functional dynamics of spinal circuits, and the underlying process of lateral inhibition, illuminate potential spinal mechanisms that could be instrumental in shaping the perception of the TGI.

Further, our observation that spatial factors influence the TGI suggests possible neuroanatomical and functional asymmetries in thermosensory and pain mechanisms (Fig. 6). Notably the increase in the intensity of the TGI when afferents map onto caudal cold and rostral warm segments in the spinal cord could mean a greater number of ascending fibres carrying thermosensory information in the Lissauer’s tract, or an uneven distribution of ascending and descending collaterals of propriospinal neurons (Anatomical Hypotheses). Additionally, there could be differing effects of inter-segmental inhibition between cold and warm projections along the rostrocaudal axis that lead to the effects of segmental location on TGI perception (Functional Hypothesis). This inter-segmental inhibition might reveal a directional pattern, such as stronger inhibitory signals from higher to lower spinal segments that are specific to cold, which are weaker in the opposite direction. Further research is needed to illuminate the specific anatomical and functional features of the spinal cord that influence the changes to thermosensory and painful sensations associated with the TGI identified in this study.

## Conclusion

Illusions in the thermo-nociceptive system can be leveraged to improve our understanding of mechanisms contributing to pain perception. Here, we presented results supporting the notion that the spinal cord plays a crucial role in the integration and processing of thermal information, contributing to the perception of both thermosensory enhancement and illusory pain within the TGI. Therefore, the initial mechanisms that lead to TGI percepts are likely to take place in the spinal cord. Additionally, we reported findings on directional inter-segmental effects in spinal integration underlying TGI, particularly when cold sensory afferents terminated in more caudal spinal segments than warm. Further research is needed to elucidate the neuroanatomical and functional properties of the spinal cord, as well as the intricate interplay between supraspinal and spinal processes that give rise to both the synthetic heat and burning sensations of the TGI.

## Authors contributions

Author contributions listed alphabetically according to CRediT taxonomy:

- Conceptualization: DEC, JFE, FF, PH, AGM.
- Data curation: JFE, AGM.
- Formal analysis: JFE, FF, AGM.
- Funding acquisition: FF.
- Investigation: DEC, JFE, AGM.
- Methodology: DEC, JFE, FF, AGM, AVS.
- Project administration: FF, AGM.
- Resources: FF, AGM.
- Software: JFE, FF, AGM.
- Supervision: FF, AGM.
- Visualization: JFE, FF, AGM.
- Writing – original draft: FF, AGM.
- Writing – review & editing: JFE, FF, PH, AGM, AVS.

## Supporting information

Suppelementary Materials

## Acknowledgements

We thank Małgorzata Basińska, Patrik Molnár, Sára A. Szabó for their help with participant recruitment and data collection. We also thank Michael Clements for the collection of pilot data that was instrumental in refining the experimental design. This study was supported by a European Research Council Starting Grant (ERC-2020-StG-948838).

