## Supplementary material for "Disentangling the spinal mechanisms of illusory heat and burning sensations in the Thermal Grill Illusion": Suppelementary Materials

### Supplementary materials

Tables of main effects and interactions for each sensory quality (cold, warm, burning) for experiment 1 and experiment 2.

#### Cold ratings

| Exp. 1: Cold reference on cold ratings | | | | |
|---|---|---|---|---|
| beta~manipulation * cold_probe + trial_n + (1 \| ID) + (1 \| order), family = beta(link = logit) | | | | |
| | β | SE | Z | p |
| Intercept | -0.88 | 0.17 | -5.2 | 2.5e-07 |
| TGI-CNT | -0.19 | 0.027 | -7 | 1.9e-12 |
| Within-Across | -0.15 | 0.037 | -4 | 6e-05 |
| Caudal-Rostral | -0.15 | 0.052 | -2.9 | 0.0038 |
| Proximal-Distal | -0.18 | 0.053 | -3.3 | 0.00094 |
| Trial | 0.0074 | 0.00099 | 7.5 | 6.3e-14 |
| TGI-CNT * Within-CNT | -0.15 | 0.054 | -2.7 | 0.007 |
| TGI-CNT * Caudal-Rostral | -0.099 | 0.075 | -1.3 | 0.19 |
| TGI-CNT * Proximal-Distal | -0.082 | 0.079 | -1 | 0.3 |

| Exp. 2: Warm reference on cold ratings | | | | |
|---|---|---|---|---|
| beta~manipulation * cold_probe + trial_n + (1 \| ID) + (1 \| order), family = beta(link = logit) | | | | |
| | β | SE | Z | p |
| Intercept | -2.2 | 0.15 | -15 | 4.6e-50 |
| TGI-CNT | 1.1 | 0.046 | 23 | 3.9e-115 |
| Within-Across | 0.013 | 0.063 | 0.22 | 0.83 |
| Caudal-Rostral | -0.01 | 0.089 | -0.11 | 0.91 |
| Proximal-Distal | -0.18 | 0.088 | -2 | 0.044 |

| Exp. 1: Cold reference on cold ratings | | | | |
|---|---|---|---|---|
| beta~manipulation * cold_probe + trial_n + (1 \| ID) + (1 \| order), family = beta(link = logit) | | | | |
| | β | SE | Z | p |
| Trial | 0.0099 | 0.0013 | 7.6 | 3.8e-14 |
| TGI-CNT * Within-CNT | 0.082 | 0.076 | 1.1 | 0.28 |
| TGI-CNT * Caudal-Rostral | -0.23 | 0.11 | -2.1 | 0.033 |
| TGI-CNT * Proximal-Distal | 0.1 | 0.11 | 0.96 | 0.34 |

### Warm ratings

| Exp.1: Cold reference on warm ratings | | | | |
|---|---|---|---|---|
| beta~manipulation * cold_probe + trial_n + (1 \| ID) + (1 \| order), family = beta(link = logit) | | | | |
| | β | SE | Z | p |
| Intercept | -2 | 0.12 | -17 | 4.5e-63 |
| TGI-CNT | 0.74 | 0.048 | 15 | 1.1e-53 |
| Within-Across | 0.26 | 0.075 | 3.5 | 0.00053 |
| Caudal-Rostral | 0.21 | 0.11 | 2 | 0.048 |
| Proximal-Distal | -0.078 | 0.1 | -0.78 | 0.44 |
| Trial | 0.0018 | 0.0016 | 1.1 | 0.26 |
| TGI-CNT * Within-CNT | -0.026 | 0.092 | -0.29 | 0.77 |
| TGI-CNT * Caudal-Rostral | 0.062 | 0.14 | 0.46 | 0.65 |
| TGI-CNT * Proximal-Distal | 0.19 | 0.13 | 1.5 | 0.14 |

| Exp. 2: Warm reference on warm ratings | | | | |
|---|---|---|---|---|
| beta~manipulation * cold_probe + trial_n + (1 \| ID) + (1 \| order), family = beta(link = logit) | | | | |
| | β | SE | Z | p |
| Intercept | -0.52 | 0.15 | -3.4 | 0.00059 |
| TGI-CNT | 0.035 | 0.029 | 1.2 | 0.23 |
| Within-Across | -0.039 | 0.036 | -1.1 | 0.28 |
| Caudal-Rostral | 0.19 | 0.051 | 3.7 | 0.00018 |
| Proximal-Distal | -0.042 | 0.051 | -0.81 | 0.42 |
| Trial | 0.0037 | 0.00094 | 3.9 | 9.1e-05 |
| TGI-CNT * Within-CNT | 0.074 | 0.052 | 1.4 | 0.16 |
| TGI-CNT * Caudal-Rostral | 0.006 | 0.073 | 0.081 | 0.94 |
| TGI-CNT * Proximal-Distal | 0.0035 | 0.074 | 0.047 | 0.96 |

### Burning ratings

| Exp. 1: Cold reference on burning ratings | | | | |
|---|---|---|---|---|
| beta~manipulation * cold_probe + trial_n + (1 \| ID) + (1 \| order), family = beta(link = logit) | | | | |
| | β | SE | Z | p |
| Intercept | -1.4 | 0.16 | -8.6 | 6.8e-18 |
| TGI-CNT | 0.39 | 0.035 | 11 | 4.5e-29 |
| Within-Across | -0.044 | 0.051 | -0.86 | 0.39 |
| Caudal-Rostral | -0.12 | 0.072 | -1.6 | 0.11 |
| Proximal-Distal | -0.17 | 0.073 | -2.3 | 0.021 |
| Trial | 0.0046 | 0.0013 | 3.7 | 0.00021 |
| TGI-CNT * Within-CNT | 0.1 | 0.069 | 1.5 | 0.15 |
| TGI-CNT * Caudal-Rostral | 0.06 | 0.097 | 0.61 | 0.54 |
| TGI-CNT * Proximal-Distal | -0.087 | 0.098 | -0.89 | 0.37 |

| Exp. 2: Warm reference on burning ratings | | | | |
|---|---|---|---|---|
| beta~manipulation * cold_probe + trial_n + (1 \| ID) + (1 \| order), family = beta(link = logit) | | | | |
| | β | SE | Z | p |
| Intercept | -1.2 | 0.17 | -7 | 2.7e-12 |
| TGI-CNT | 0.37 | 0.034 | 11 | 4.5e-27 |
| Within-Across | -0.068 | 0.043 | -1.6 | 0.11 |
| Caudal-Rostral | 0.17 | 0.061 | 2.8 | 0.0054 |
| Proximal-Distal | -0.003 | 0.062 | -0.049 | 0.96 |
| Trial | 0.0075 | 0.0011 | 7 | 3.2e-12 |
| TGI-CNT * Within-CNT | 0.21 | 0.059 | 3.6 | 0.00036 |
| TGI-CNT * Caudal-Rostral | -0.14 | 0.084 | -1.7 | 0.096 |
| TGI-CNT * Proximal-Distal | -0.058 | 0.084 | -0.69 | 0.49 |

### Temperatures by participant

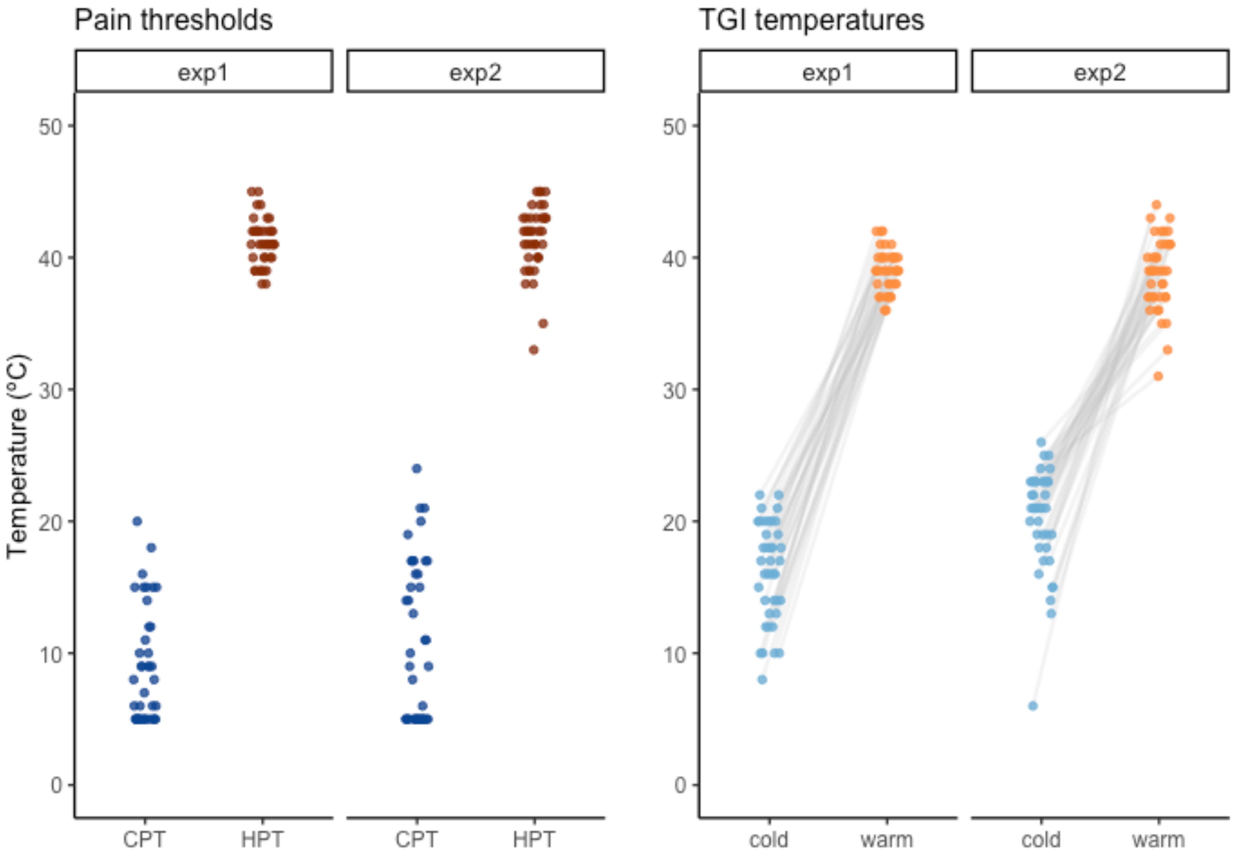


*Figure S1: Pain thresholds (left) and TGI temperatures (right) for each participant in experiment 1 and experiment 2. As we did not go below 5ºC or above 45ºC for the TGI stimuli, CPT and HPT were capped at 5ºC and 45ºC respectively.*
